# Automatic Analysis of Bees’ Waggle Dance

**DOI:** 10.1101/2020.11.21.354019

**Authors:** Jordan Reece, Margaret Couvillon, Christoph Grüter, Francis Ratnieks, Constantino Carlos Reyes-Aldasoro

## Abstract

This work describe an algorithm for the automatic analysis of the waggle dance of honeybees. The algorithm analyses a video of a beehive with 13,624 frames, acquired at 25 frames/second. The algorithm employs the following traditional image processing steps: *conversion to grayscale, low pass filtering, background subtraction, thresholding, tracking and clustering* to detect run of bees that perform waggle dances. The algorithm detected 44,530 waggle events, i.e. one bee waggling in one time frame, which were then clustered into 511 *waggle runs*. Most of these were concentrated in one section of the hive. The accuracy of the tracking was 90% and a series of metrics like intra-dance variation in angle and duration were found to be consistent with literature. Whilst this algorithm was tested on a single video, the ideas and steps, which are simple as compared with Machine and Deep Learning techniques, should be attractive for researchers in this field who are not specialists in more complex techniques.

## I. Introduction

The honeybee waggle dance is one of the most complex forms of communication in the animal kingdom and can be good indicator of seasonal foraging challenges [1]. Upon finding a profitable food source, the bee returns to its hive and performs a waggle dance to convey its location to her hive-mates. The dance consists of a figure-of-eight motion known as the ‘return phase’, followed by a ‘waggle run’ in the eight’s centre, where the bee shakes its body from side to side while moving forward in a straight line [2]. These two phases comprise a circuit, which can be repeated many times, as few as 1 as many as 100 [3]. The dance can recruit bees to the newly discovered source, thus maximising the profitability of the recent finding and reducing the need for scouting for new food sources, which can be costly for a hive.

From the waggle run alone, the direction, distance and profitability of the food source can be conveyed to the other bees in the hive. Bees, like many other insects, use the sun for navigation purposes by comparing the suns position to various landmarks [4]. The direction of the food source is conveyed by the bees’ mean orientation throughout the waggle run relative to the upward direction on the vertical comb [5]. The distance flown to the food source is communicated by the duration of the waggle run [6]. Profitability requires information on the dance as a whole, with duration of the entire dance (several waggle runs) and the number of waggle runs per dance being particularly good indicators [7].

Although most labs will now use digital recordings, they still mostly rely on manual measurements, which can be time consuming [3], [8]. Automated methods of recording the honeybee dance language can free researchers from hundreds of hours of recording and help filter out particular dance behaviours for more nuanced research questions. Other groups have already attempted to automate dance language recording with some success [8], [9]. Current limitations centre around working with lower quality footage, the level of noise in the hive and calculation errors.

In this work, we describe an algorithm that can automatically detect waggle runs, record several features of the run that are useful to researchers and cluster the runs into full dances. Two datasets were analysed with varying levels of detail, a frame by frame tracking of each run and a summary of each individual run, both labelled to their respective dances. The algorithm is designed to work on lower quality footage and to require only a few parameters to optimise to a lab’s setup.

## II. Materials and Methods

### A. Materials

The footage studied was taken from an observation hive from the Laboratory of Apiculture and Social Insects (LASI) at the University of Sussex (https://www.sussex.ac.uk/lasi). The video consists of 9 minutes of 720×1280 resolution footage at 25 frames per second (fps). Whilst other studies have used higher resolution or higher frame rate videos [8], [9], regular resolution and frame rates (25-30 fps) is more accessible to all researchers and thus was the focus of this work.

### B. Methods

The algorithm was designed in Python using the OpenCV library to process the video. Results were evaluated against ground truth obtained by manual selection of a rectangle to contain each bee that waggled. The orientation was calculated with a manually drawn line from head to tail of each bee to bisect the body in halves. The algorithm consists of three main stages: (1) motion detection, (2) waggle run tracking, and (3) clustering into a waggle dance.

#### Motion Detection

The motion detection consisted of (a) the conversion of the colour images (RGB) to grayscale, (b) low pass filtering of the images with a Gaussian filter, (c) background subtraction, i.e. *frame*(*t*) − *frame*(*t* − 1), pixels that do not change (background) will be close to zero and pixels that change (bee in motion) will change and have higher intensity, and (d) thresholding to separate pixels that indicated motion. There are 2 types of movement in the hive: fast waggling movements and slower movements of displacement. Slow movements will be imply uniform variation of the bee with thin lines, whilst a waggle dance would be thicker lines more prominent in one side (tail) of the bee (Fig. 1).A waggle run in the background subtracted image appears as a disconnected contour with a larger area, compared to the hollow contour produced from normal movement (Fig. 2). Hierarchical contouring can separate disconnected contours, known as child contours, from the hollow (parent) contours of other bees. Combining hierarchical contouring with a minimum area threshold allows waggle runs to be segregated from background noise.

**Fig. 1:**
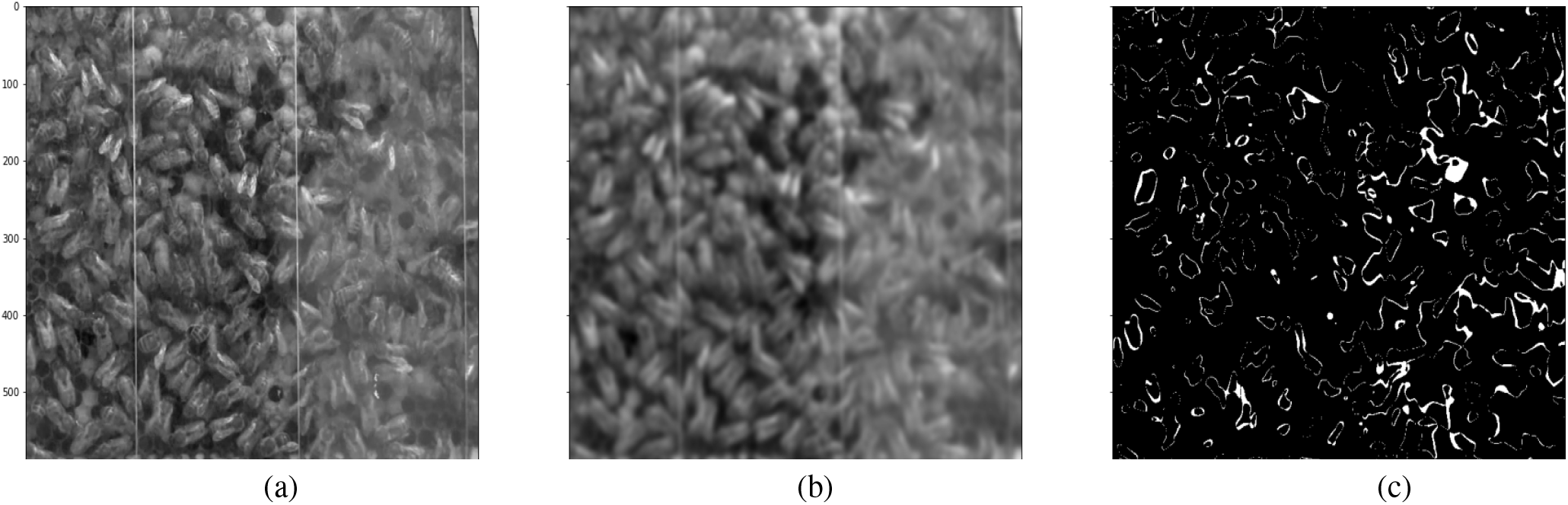
Illustration of the main pre-processing stages: (a) conversion to from RGB colour to grayscale, (b) low-pass filtering with a Gaussian blur, (c) background subtraction and thresholding.

**Fig. 2:**
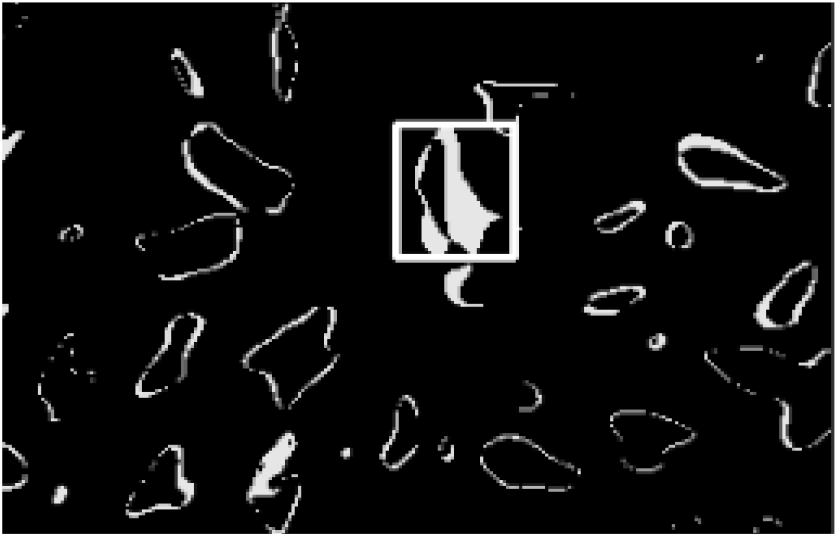
Illustration of the detection of a waggling bee, based on the thickness of edges obtained with background subtraction.

Every contour that fit the criteria for ‘waggle-like’ activity was recorded to a dataset to be clustered into waggle runs. Waggle runs consist of waggle-like activity over consecutive frames, lasting up to several seconds in approximately the same position. The Density-based spatial clustering of applications with noise (DBSCAN) algorithm [10] was used to cluster the activity into individual waggle runs using the spatial coordinates and timestamp as the features, for its effectiveness on spatiotemporal data. The algorithm finds an undefined number of clusters and therefore scales effectively without the need of additional parameters. Once the entire video has been processed, the cluster dataset was cleaned to remove possible false positives. The aim was to remove all false positives, even at the expense of true positives, which can be reinserted during the tracking phase. The resultant dataset can also include multiple datapoints for the same frame. As a bee cannot be in two places at once, these duplicates also contain false positives that need to be removed. The Euclidean distance between each point and the point at the previous frame was calculated and only the point with the shortest Euclidean distance was preserved. The datapoints in the top decile of Euclidean distance were removed, removing non-duplicates that are beyond the possible distance a bee can travel in a single frame.

#### Tracker

The effect of the strict discarding of false positive was that the returned dataset had several missing datapoints, with an average of 40% missing frames within each waggle run. To compensate for these missing frames, each bee was tracked from the first to final frame within the bounds of the cluster, using the motion detection dataset as a skeleton. The known datapoints in each cluster were used to create an interpolation function to get an estimate of the bee’s location in the missing frames. The bee was tracked throughout the missing frames using contour-based tracking. Frame by frame, a bee does not move far from its position in the previous frame. A bounding box was centred around the contour, extended beyond the size of the bee to account for the movement of the bee in the next frame. The contour was then found in the next frame and the bounding box recentred. The interpolation function was used as a reference point to prevent drift or tracking failure. A larger bounding box was created with the interpolation function’s coordinates at frame *t* as a centre. If the contour’s centre was within the interpolated bounding box, the tracker was considered successful. In the case of drift, a bounding box was created around the interpolated coordinates and the largest contour was found. The number of frames between two known datapoints are rarely more than 4 frames,with a maximum difference of 7 frames within this dataset. This method removes the need for long term tracking and therefore removes the current limitations that are associated with it. Information regarding the bee’s position and the size of its contour were recorded per frame. Orientation data was also recorded by taking the angle of the major axis of the bounding box in each frame and calculating the mean.

#### Clustering

The data for each cluster (waggle run) was then summarised in a dataset where each run is a single datapoint. Time taken, mean angle, waggle frequency, direction and distance travelled were all recorded for each waggle run. A 5s search window from the end of each run was used to find successive runs in a waggle dance. The closest run in the search window was saved as the next run in the dance sequence. The same duplicate removal strategy was applied to this dataset, removing all but the closest succeeding points when two different runs claimed the same run as the next in their sequence. The successive runs were grouped together, forming a list of runs that make up a full dance. All runs in the frame-by-frame and summarised dataset were labelled to the dance they belong to. There was a final option to save each dance as a cropped video, along with visual cues to aid researchers in manually tracking the bee.

#### Parameters

The following parameters were manually determined: values to create the binary threshold image, clustering parameters detailing the distance between each data point in the cluster and the size of the kernels used in morphological operations such as erosion and dilation.

## III. Results

A total of 13,624 video frames were analysed, 44,530 waggle events were detected and of these, 511 clusters were formed (Fig. 3). The Motion Detector results were evaluated by taking ROIS of 300 × 300 pixels over 200 frames at random locations and manually counting the number of waggle runs present in each. Over 50 manually inspected ROIs, the stage was successful with 92.8% precision and 77.2% recall. This suggests very few false positives were being detected by the algorithm, but up to 22.8% of waggle runs were missed.

**Fig. 3:**
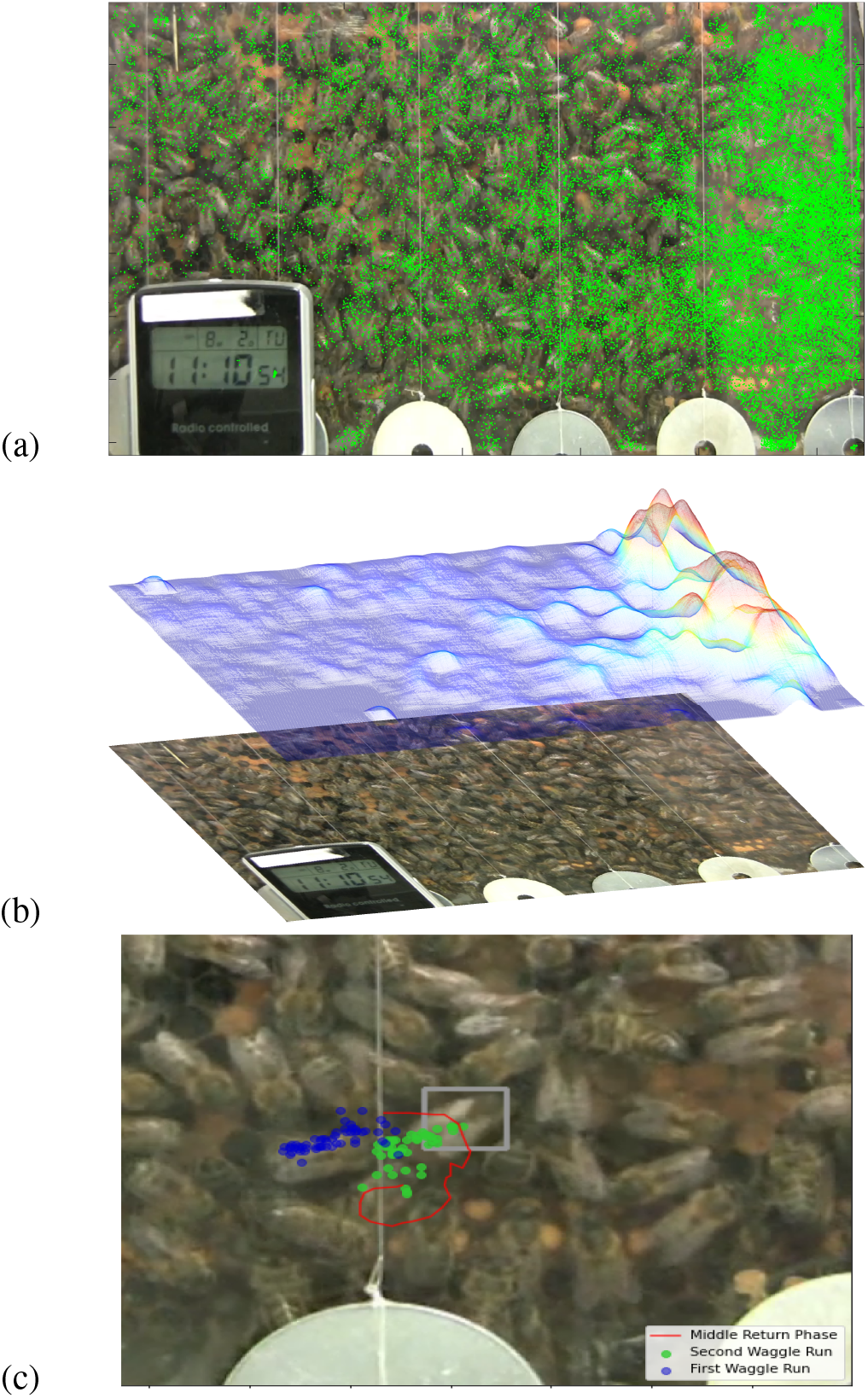
Illustration of the waggle events detected. (a) All waggle detections overlaid as a green dot. (b) A heat map indicating the frequency of the events relative to the position of the hive. (c) One event illustrated with blue and green dots for the first and second waggle runs. The bee is highlighted in the bounding box.

Tracking accuracy was measured against an overlap coefficient. The final dataset maintained 90% accuracy when tested on hundreds of frames. 79.6% of contours tested returned an overlap coefficient of 1, suggesting perfect tracking.It should be noticed that the overlap coefficient was comparing whether the area of the contour resided entirely within the bounding box and the contour was often smaller than the size of the bee. The contour overlay and ground truth bounding box displayed in Fig. 3(c) demonstrates how detection accuracy was measured. Values in the range 0 < *x* < 1 suggest an overlap between the the detected contour and a manually drawn bounding box around the bee, likely caused by under-segmentation of the image. An overlap coefficient of 0 suggests tracking failure. It should be mentioned that we were not interested in segmentation quality and thus did not measure the accuracy with intersection over union or other metrics. We were chiefly concerned with the contour of the bee being contained in the bounding box.

Orientation was measured using a similar method to the algorithm. At each frame, a line was manually drawn directly through the bee along its major axis and the angle of the line was recorded. The angles over an entire cluster were averaged to get the orientation of the run. The dataset averaged −9.3° of error from the true angle with a ±5.6° S.D. This has not improved on the −2.92° ± 7.37° error found in pre-existing methods [8].

The mean average error between the true duration in frames and recorded duration were ± 7.57 frames. Based on full duration of each dance, this averaged a 17% error rate. The median error rate stayed around −2.0 frames, suggesting that typically, the true waggle run duration was longer than that recorded in the dataset. The variation in waggle run duration was greatly increased at the end of the waggle run, with the start of the waggle run having greater accuracy.

## IV. Discussion

Reproducing the results found in the literature using the final outputted dataset demonstrated the efficacy of the algorithm here described. While the results will not be exactly the same, if findings suggested in the literature can be gleaned from the final dataset, it is evidence that the results produced are accurate.

Intra-dance variation [3], [11], [12] is another phenomenon found in the literature that is easily reproducible. Longer dances (>2 runs) were used to calculate the standard deviation from the mean angle and duration of each dance. These were then divided into 3 groups (first run, middle runs, last run) and compared in Fig. 4. The literature found a significant difference between the tail and middle runs with a p<0.001 confidence level [3]. The differences within this sample were less pronounced, with no significant difference found in the intra-dance variation with regards to waggle run duration. There was a significant difference between the angle variation (p<0.005), albeit with less confidence than the literature. Comparisons between the trends found in the literature [3] and this dataset (Fig. 4) do, however, have a lot of similarity. Duration is one aspect of the algorithm with the highest rate of error (17%) which could affect the results.

**Fig. 4:**
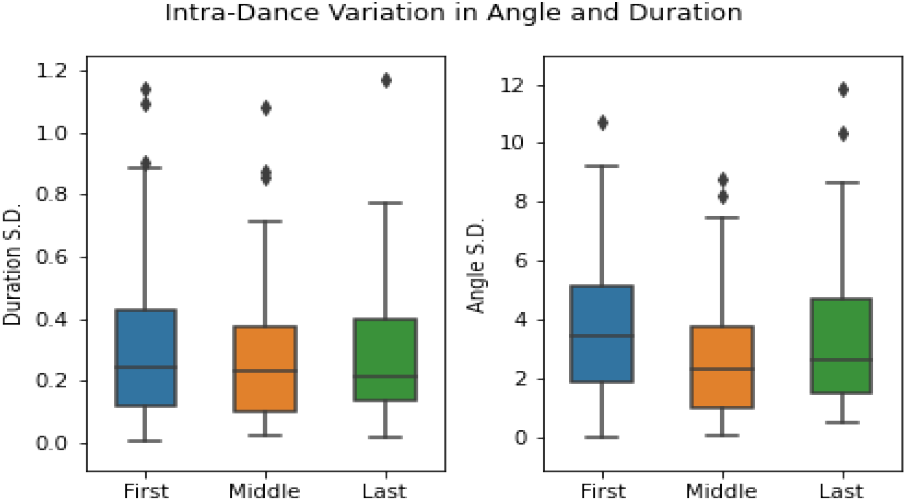
Intra-dance variability taken from the final dataset. The results mirror those found in the literature [3].

The results show that the methods perform with a high level of accuracy. The final dataset can mirror some of the results found in the literature using manual methods that can take several minutes to decode per dance [3]. With ~ 500 waggle runs recorded from a 10-minute video, this method can save researchers hours of time, with only an initial parameter calibration required on any one setup. Reducing the time spent measuring bees’ movements and increasing the quantity of data available to researchers is of benefit to the field.

The detection and tracking accuracy both perform to a high standard detecting 77% of all waggle runs and accurately following the bee 90% of the time. Increasing the motion detection sensitivity parametrically will increase the frequency of false positives but may be of use to researchers. An additional postprocessing step can filter out false positives by comparing the waggle frequency of true and false positives. False positives are likely to have a frequency greatly below the 15 Hz prescribed in the literature.

The orientation error is likely caused by the tracker primarily tracking the wings of the bee instead of its body. As bees splay their wings during the waggle run, this can lead to a slightly different recording when compared to the true orientation. This can be adjusted for by reducing the minimum threshold value within the bounding box until the resulting contour is nearer the full size of the bee. The error in waggle run duration is a systematic error caused by the hard bounds created from the initial DBSCAN clustering. Extending tracking beyond the bounds set by the cluster can verify whether the cluster stopped the recording prematurely. Additional postprocessing can compare the similarity of the bee’s movements post-cluster to the movement during the waggle run. Whilst we only ran on one video, we think that the nature of the steps of the pipeline can generalise to videos with similar quality and conditions.

In this work we described an automatic algorithm that detected bees that were performing a waggle dance. The algorithm allows the extraction of metrics frequently found in the literature. The results were compared against ground truth data and the algorithm shows promise of aiding researchers in this labour-intensive task. Future work will focus on improving the orientation and duration error, the means by which have been outlined within this paper.

The code is available “as is” through GitHub: *https://github.com/Jreece18/WaggleDanceTracker*.

